# Population demographic history and evolutionary rescue: influence of a bottleneck event

**DOI:** 10.1101/2023.01.11.523672

**Authors:** Laure Olazcuaga, Beatrice Lincke, Sarah Delacey, Lily F. Durkee, Brett A. Melbourne, Ruth A. Hufbauer

**Affiliations:** Department of Agricultural Biology, Colorado State University, Fort Collins, Colorado 80523, USA; Graduate Degree Program in Ecology, Colorado State University, Fort Collins, Colorado 80523, USA; Department of Ecology & Evolutionary Biology, University of Colorado, Boulder, Colorado 80309, USA

**Keywords:** adaptation, genetic variation, inbreeding depression, heterozygosity, extinction, *Tribolium castaneum*

## Abstract

Rapid environmental change presents a significant challenge to the persistence of natural populations. Rapid adaptation that increases population growth, enabling populations that declined following severe environmental change to grow and avoid extinction, is called evolutionary rescue. Numerous studies have shown that evolutionary rescue can indeed prevent extinction. Here, we extend those results by considering the demographic history of populations. To evaluate how demographic history influences evolutionary rescue, we created 80 populations of red flour beetle, *Tribolium castaneum*, with three classes of demographic history: diverse populations that did not experience a bottleneck, and populations that experienced either an intermediate or a strong bottleneck. We subjected these populations to a new and challenging environment for six discrete generations and tracked extinction and population size. Populations that did not experience a bottleneck in their demographic history avoided extinction entirely, while more than 20% of populations that experienced an intermediate or strong bottleneck went extinct. Similarly, among the extant populations at the end of the experiment, adaptation increased the growth rate in the novel environment the most for populations that had not experienced a bottleneck in their history. Taken together, these results highlight the importance of considering the demographic history of populations to make useful and effective conservation decisions and management strategies for populations experiencing environmental change that pushes them toward extinction.

## Introduction

In the Anthropocene, populations experience frequent and intense changes to their environments (Bellard *et al*., 2012). If environmental change is severe enough, population growth may slow to the point that population size starts to decline. If dispersal capabilities are limited or suitable environments are no longer available (Schloss *et al*., 2012), populations must adapt to challenging environments or risk extinction (Smith *et al*., 1989; Holt, 1990). Adaptation that is rapid enough to reverse population decline due to a challenging environment and prevent an otherwise inevitable extinction is called evolutionary rescue (Gomulkiewicz & Holt, 1995; Carlson *et al*., 2014). This process is characterized by a U-shaped curve in population size through time: initially, population size decreases due to maladaptation to the environment, then population size increases as adaptive alleles spread through the population (Gomulkiewicz & Holt, 1995; Orr & Unckless, 2014).

Theory and experiments with model organisms show that evolutionary rescue can avert extinction. Theory has demonstrated that the probability of evolutionary rescue increases with genetic variation (Gomulkiewicz & Holt, 1995; Barrett & Schluter, 2008) and suggests that it decreases with mutational load and inbreeding depression (Barrett & Kohn, 1991; Ellstrand & Elam, 1993). Despite confirmation of these effects in experiments with model organisms (Bijlsma *et al*., 2010; Agashe *et al*., 2011; Lachapelle & Bell, 2012; Ramsayer *et al*., 2013; Hufbauer *et al*., 2015), evidence of evolutionary rescue in nature remains scarce, suggesting that the conditions that allow evolutionary rescue to occur may be limited (Carlson *et al*., 2014). One condition that is particularly important to examine experimentally is the demographic history of populations. In the wild, populations are frequently subjected to various stressful events that can lead to a temporary reduction in size, such as hunting or habitat destruction. These bottlenecks in population size (Frankham *et al*., 1999) can affect genetic variation, mutational load, and inbreeding levels in the populations that experience them (hereafter bottlenecked populations). Such bottleneck events, whether rare extreme events or repeated less-severe restrictions in population size, can reduce the future adaptive potential of populations (Nei *et al*., 1975; Bouzat, 2010) and thus the probability of evolutionary rescue.

Bottlenecks influence the ability of populations to adapt to a future challenging environment in two main ways (O’Brien & Evermann, 1988; Frankham *et al*., 1999; Reusch & Wood, 2007). First, standing genetic variation is lost by genetic drift during bottlenecks due to an increase of stochastic events associated with small population size (Hedrick, 2005). This reduction of genetic diversity decreases the evolutionary potential of populations since potential future adaptive alleles are lost (Futuyma, 1986). In general, adequate genome-wide genetic variation is important for the conservation of natural populations (Kardos *et al*., 2021). In addition, when populations are small, selection is weak relative to genetic drift. When this occurs, there can be an increase in the frequency and fixation of deleterious alleles (Lynch *et al*., 1995), leading to populations with lower growth rates (average number of offspring per individual) than populations that have not experienced a bottleneck.

However, given the stochastic way in which alleles go to fixation or are lost, by chance, some beneficial alleles may become fixed, potentially leading some bottlenecked populations to have higher growth rates than non-bottlenecked populations (Bouzat, 2010). Thus, in addition to a lower growth rate, greater variation in growth rate is expected across different bottlenecked populations compared to non-bottleneck populations.

The second main way that bottlenecks can influence the ability of populations to adapt is through inbreeding. After bottleneck events, inbreeding increases due to increased homozygosity of alleles common by descent (Wright, 1977; Ralls *et al*., 1979; Frankham *et al*., 1999; Kirkpatrick & Jarne, 2000; Fox *et al*., 2008; Charlesworth & Willis, 2009; Hedrick & Fredrickson, 2010). When homozygosity increases at loci with deleterious alleles, inbreeding leads to inbreeding depression (i.e., a reduction in absolute fitness of inbred individuals). Inbreeding depression increases the risk of extinction for the population by reducing its growth rate (Charlesworth & Charlesworth, 1987; Frankham, 2005a, 2005b; O’Grady *et al*., 2006). However, inbreeding in populations can also lead to purging of deleterious alleles, removing them from the population and increasing growth rate of the population (Crnokrak & Barrett, 2002; Glémin, 2007; Facon *et al*., 2011). Thus, as for genetic drift, the effects of inbreeding due to bottlenecks can increase the variation in growth rate among populations.

Despite the strong evidence for a positive effect of genetic variation (Hedrick, 2005) and a negative effect of inbreeding depression (Frankham *et al*., 1999) on the rapid adaptation that is central to evolutionary rescue, the consequences of past bottleneck events on adaptation have only recently begun to be explored.

Experiments using bacteria demonstrate that the adaptive pathways followed by populations under stress depend on whether bottlenecks are weak or intense (Vogwill *et al*., 2016; Garoff *et al*., 2020; Mahrt *et al*., 2021), supporting the idea that demographic history will influence the probability of evolutionary rescue. In those experiments, with large bacterial populations, *de novo* mutations generated genetic diversity. However, in natural diploid populations with sexual reproduction, adaptation relies mostly on standing genetic variation, which should result in very different adaptation dynamics. Furthermore, in a recent study, Klerks *et al*., (2019) evaluated how population demographic history (bottlenecked or not) influenced the response to selection for increased heat tolerance in experimental populations of a diploid, sexually reproducing eucaryote: the least killifish *Heterandria formosa*. They found that bottlenecked populations had a 50% slower response to selection than normal populations. This suggests that bottlenecks play an important role in the outcomes of evolutionary rescue. However, monitoring population dynamics was outside the scope of Klerks *et al*., (2019) who studied adaptation in experimental populations of fixed size, and so it is not clear if the U-shaped curve characteristic of evolutionary rescue would have been observed had population size been allowed to vary. Studying population dynamics is, however, necessary to understand the minimum population size that populations will reach without going extinct or how long it will take for populations of a given size to recover. Given the role of bottlenecks in reducing genetic variation and increasing inbreeding depression in diploid eukaryotes that are often of conservation concern (Frankham *et al*., 1999; Reusch & Wood, 2007), it is important to examine the effects of bottlenecks on evolutionary rescue experimentally.

Here, we studied how bottleneck events influence the process of evolutionary rescue of populations in a severely challenging environment. Because bottlenecks in nature vary in intensity, we created populations that had experienced no bottleneck in their demographic history, an intermediate bottleneck, or a strong bottleneck, and monitored population dynamics in a challenging environment. By using *Tribolium castaneum* to model a diploid, semelparous organism, we evaluated how the probability of extinction depended upon population demographic history, and whether surviving populations exhibited the U-shaped population trajectory characteristic of evolutionary rescue, and how that depended upon population demographic history. Finally, we quantified adaptation to the new environment at the end of the experiment by measuring the growth rate of all rescued populations and evaluating the evolution of the rate of development, a life history trait that could be involved in adaptation (Berner *et al*., 2004).

## Methods

### Model system

We used *T. castaneum*, the red flour beetle, to test the effects of demographic history on evolutionary rescue. *Tribolium castaneum* populations were kept in 4 cm by 4 cm by 6 cm acrylic containers with 20 g of a standard medium (95% wheat flour and 5% brewer’s yeast), which served as both habitat and a food source. We will henceforth refer to a container and medium together as a patch. Multiple patches were used to maintain each distinct laboratory population, and single patches were used in growth assays. During laboratory maintenance, the life cycle of *T. castaneum* was constrained to mimic that of a seasonally breeding organism with non-overlapping generations and a discrete dispersal phase (Melbourne & Hastings, 2008), a life history found commonly among plants and animals. Each generation, adult beetles from all the patches were mixed. Then, we placed them in groups of 50 individuals per patch with fresh standard medium for 24 hours to reproduce and start the next generation. At the end of the reproduction phase, adults were removed from the patches and the eggs were left to mature into adults for an additional 34 days for a total of 35 days per generation. During laboratory maintenance and the experiments, beetles were kept in incubators at 31°C with 54 ± 14% relative humidity. Several incubators were used, and patches were randomized among and within incubators once each week to prevent systematic effects of incubation conditions.

### Creating populations with different demographic histories

A large diverse population was initiated from the multiple crosses of different stock strains (“GA1”, “Lab-S”, “Z2”, “Z4” and “Estill” from USDA stock maintenance). These strains were originally all from samples from the US southwest, to minimize potential genetic or cytoplasmic incompatibilities between strains. All possible pairwise crosses indicated no reduction in fitness between any given pair (Durkee et al. 2023). This diverse population was maintained and well mixed for approximately ten generations at a population size of 10,000 individuals. This population was divided into four subsets spaced one week apart (hereafter “temporal blocks”), with gene flow between the different temporal blocks. From this large and diverse population, we created 60 independent replicate populations experiencing bottlenecks in population size of two different intensities over two generations. During the first generation of the bottleneck event, all 60 populations were reduced to a size of two individuals. During the second generation of the bottleneck event, 24 populations were allowed to triple in size to six individuals to initiate the next generation (hereafter called “intermediate bottleneck”) and 36 populations were not allowed to grow, and experienced a second bottleneck of two individuals, to initiate the next generation (hereafter called “strong bottleneck”). Contrary to our expectations, the success rate of bottleneck population creation was the same regardless of bottleneck intensity, thus we successfully created more replicates for strong bottleneck populations than for intermediate bottleneck populations. Then, these 60 populations were maintained for six additional generations in the laboratory to increase population size prior to starting the experiment (Fig. 1). In contrast to the intermediate and strong bottlenecked populations, the diverse population was maintained as a single large population during those six generations to maintain high genetic diversity and was used to create independent replicate populations at the beginning of the experiment (hereafter called “diverse populations”). During the six generations prior to the initiation of the experiment, the population size of each bottlenecked population was recorded so that the individual population demographic history within each bottleneck category was known.

**Figure 1.**
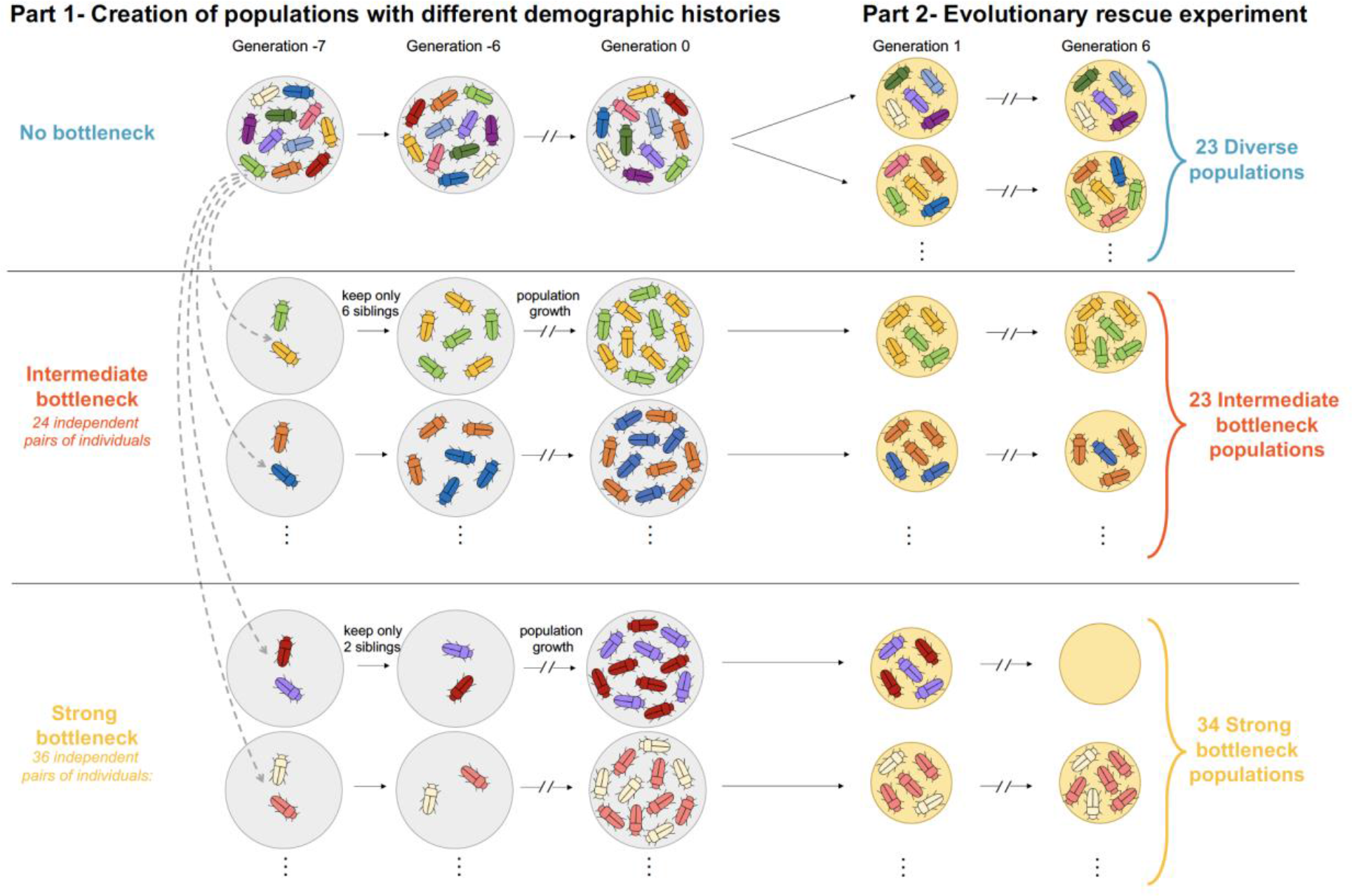
Experimental design. The bottlenecked populations experienced a bottleneck event seven and eight generations before the beginning of the experiment (generations represented by negative values G -7 and G -6). During these eight preliminary generations (Part 1, left), the populations were maintained on a rich environment (grey circles). During the experiment (Part 2, right), the populations evolved on a stressful medium mainly composed of corn flour (yellow circles).

### Quantifying the demographic history

By tracking the population size of each population each generation, we know their full demographic history since the bottleneck. We calculate the expected heterozygosity remaining in each population relative to the source population as an index that takes into account the variation in population size through time. Following Allendorf (1986), the expected heterozygosity at generation *t* (*He*_*t*_), is a function of expected heterozygosity and population size in generation *t*-1:

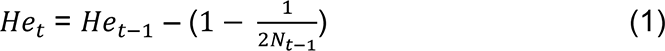

Where *He*_*t*_ is the expected heterozygosity at the previous generation *t*-1 and *N*_*t*−1_ is the population size at the previous generation *t*-1. To calculate the proportion of expected heterozygosity remaining, expected heterozygosity of the source population at the time of the bottleneck is defined as 1, that is *He*_0_ = 1. The most accurate way to calculate this is using effective population size, *N*e (i.e., the size of an ideal population that would lose genetic variation at the same rate as the focal population; Allendorf 1986). We used census population size (*N*), as that is what we had available. In the case of *T. castaneum* these values are correlated (e.g., *N*e/*N* around 0.9 for *N* = 100 individuals, Pray et al. 1996; Wade 1980). Because *N* is slightly larger than *N*e, our calculation is biased in a conservative way, estimating smaller losses of heterozygosity than may have actually occurred. This calculation enables an additional evaluation of the consequences of our experimentally imposed bottlenecks on evolutionary rescue. It also provides a way to compare the strength of the bottlenecks imposed on our experimental populations with other bottleneck situations. For example, the intermediate bottleneck is comparable to a population remaining at a size of 50 individuals for about 37 generations, and the strong bottleneck is comparable to a population remaining at a size of 50 individuals for about 57 generations.

### Evolutionary rescue experiment

To test the effect of demographic history on evolutionary rescue, we compared the probability of rescue of diverse populations to that of bottlenecked populations. In total, we compared the evolution of 36 diverse populations, 24 intermediate bottleneck populations, and 36 strong bottleneck populations distributed over three temporal blocks. To start the experiment, 100 individuals from each population were housed in two patches connected via 2mm holes drilled in the sides. We challenged these experimental populations over six discrete generations with a stressful medium that consisted of 97% corn flour, 2.85% wheat flour, and 0.15% brewer’s yeast, to represent a resource-limited environment similar to Hufbauer *et al*. (2015). Each generation, the adults in each population were censused completely, then the individuals were mixed and put back equally in the two connected patches with fresh challenging media for the 5-week life cycle as described above. For each population, the proportion of expected heterozygosity was computed each generation (Eq. 1) until the end of the experiment or the extinction of the population. A population was considered extinct when the population size was 0 or 1 at census. At the end of the experiment, the growth rate of each population was measured as the number of offspring produced per 30 adults (replicated as much as possible depending on population size). This number of adults was chosen to enable measurements of the growth rate of all the extant populations, even those with small populations, (so as not to bias our findings by excluding smaller, potentially less adapted populations). To evaluate potential differences in development time among the populations that survived to the end of the experiment (hereafter, extant populations), the composition of different developmental stages was enumerated after 4 and 5 weeks of development by counting the number of larvae, pupae, and adults in each patch.

Due to contamination with a protozoan parasite, 12 populations were excluded from the experiment (9, 1 and 2 populations for diverse, intermediate bottleneck, and strong bottleneck populations respectively). With these exclusions, 23 diverse populations, 23 intermediate bottleneck populations and 34 strong bottleneck populations were evaluated in the statistical analysis (Fig. 1).

### Statistical analysis

All statistical models were fitted in R version 3.4.2 (R Core Team, 2014). For all models, we estimated the significance of interactions and main effects by comparing a full model (i.e., with model terms of the same order) to a reduced model without the interaction or main effect of interest using a Likelihood Ratio Test (LRT). By using likelihood ratio tests to eliminate non-significant effects, we identified the most parsimonious model (hereafter “best-fit model”) for each dataset.

To understand how the probability of extinction in the challenging environment was influenced by demographic history, we used generalized linear models for the probability of extinction and time to extinction. Since none of the diverse populations went extinct during the experiment, the parameter estimate for this treatment in standard logistic regression and survival analysis would be infinite and estimates of uncertainty and hypothesis tests would be invalid, a phenomenon known as complete separation (Albert & Anderson, 1984). To manage this issue, we separately analyzed (1) the probability of extinction at the last generation, (2) the time to extinction of the populations that went extinct during the experiment, (3) survival of populations over time, in each case using approaches compatible with complete separation. We modeled the probability of extinction as a Bernoulli distribution with logit link fitted with penalized likelihood and Firth’s correction using the package *logistf* (Heinze *et al*., 2022). Explanatory variables in the model were demographic history (a categorical variable with three levels: “diverse”, “intermediate bottleneck” and “strong bottleneck”) and temporal block (a categorical variable with three levels), both as fixed effects (Table 1). We modeled time to extinction (number of generations) as a Poisson distribution with log link after confirming that overdispersion was absent. Explanatory variables in the model were the categorical demographic history and the temporal block, again as fixed effects (Table 1). We modeled survival over time with a survival analysis using a log-rank statistic to compare the survival dynamics across groups using the package *survival* (Therneau, 2018). The explanatory variable in the model was demographic history (a categorical variable with three levels: “diverse”, “intermediate bottleneck” and “strong bottleneck”) as a fixed effect (Fig. S1). Due to the problem of complete separation, a Cox-PH model was not fitted, as it would result in degenerate estimates of the probability of survival or extinction or time to extinction.

**Table 1.** Likelihood ratio test table for each model comparison analyzing the data measured during and at the end of the experiment. df: degrees of freedom. D: deviance computed from the likelihood or the penalized likelihood (see methods section). △df: change in degrees of freedom between the two models. △D: change in deviance between the two models (i.e., statistic of the LRT following a X2 distribution). p-value: p-value of the LRT.

| | df | D | $\Delta df$ | $\Delta D$ | *p*-value |
| --- | --- | --- | --- | --- | --- |
| **Probability of extinction (effect of demographic history)** |  |  |  |  |  |
| Extinction_probability ~ Block | 3 | 21.29 |  |  |  |
| Extinction_probability ~ Demographic_history + Block | 5 | 11.90 | 2 | 9.39 | 0.009 |
| **Time of extinction (effect of demographic history)** |  |  |  |  |  |
| Time_extinction ~ Block | 3 | 0.95 |  |  |  |
| Time_extinction ~ Demographic_history + Block | 4 | 0.75 | 1 | 0.21 | 0.65 |
| **Dynamic of the population (generation-demographic history interaction)** |  |  |  |  |  |
| Population_size ~ Demographic_history + Block + s(Generation) + s(Population, bs="re") | 8 | 1977.92 |  |  |  |
| Population_size ~ Demographic_history + Block + s(Generation) + s(Population, bs="re") + s(Generation, by = Demographic_history) | 14 | 1974.92 | 6 | 3.00 | 0.42 |
| **Dynamic of the population (effect of generation)** |  |  |  |  |  |
| Population_size ~ Demographic_history + Block + s(Population, bs="re") | 6 | 2072.84 |  |  |  |
| Population_size ~ Demographic_history + Block + s(Generation) + s(Population, bs="re") | 8 | 1977.92 | 2 | 94.92 | < 0.001 |
| **Dynamic of the population (effect of demographic history)** |  |  |  |  |  |
| Population_size ~ Block + s(Generation) + s(Population, bs="re") | 6 | 1993.35 |  |  |  |
| Population_size ~ Demographic_history + Block + s(Generation) + s(Population, bs="re") | 8 | 1977.92 | 2 | 15.43 | < 0.001 |

A U-shaped curve in population size characterizes evolutionary rescue. To evaluate if and how the shape of rescue curves is influenced by demographic history, we focused on populations extant at the end of the experiment and analyzed size through time using a Generalized Additive Mixed Model (GAMM; Wood, 2017) with a negative binomial distribution to account for overdispersion using the package *mgcv* (Wood, 2022). To allow for the nonlinear relationship, we modeled population size (starting at generation 2, the peak in abundance before the decline) as a smooth function of generation (a continuous variable from 2 to 6) using a thin plate regression spline (Wood, 2017). Explanatory variables included categorical demographic history, the interaction between generation and demographic history (thus allowing the shape of the curve to vary by demographic history) and temporal block (three levels) as fixed effects and population identity as a random effect. This model was compared with a reduced model (Table 1) using the *compareML()* function of the package *itsadug* (Jacolien et al. 2022).

In addition, to quantify the variability of responses and visualize whether outcomes of populations that experienced a bottleneck event are more variable than populations that did not, we computed the coefficients of variation in population size across the three demographic histories across all generations for the extant populations. We used the package *DescTools* to compute coefficients of variation and their 95% confidence intervals (Signorell *et al*., 2022). Considering the coefficient of variation instead of the variance allows us to compare variation while controlling for mean population size.

To test the influence of demographic history on adaptation, we evaluated growth rate under standard density and the distribution of developmental stages of extant populations. The growth rate (i.e., number of offspring divided by the number of parents) is commonly analyzed with a lognormal transformation. Due to the presence of 0 in our dataset, this direct transformation was not possible. To consider appropriate transformations, we checked the distribution of residuals from growth rate analyses with either a log transformation of growth rate + 1, a square root transformation, or without transformation. This preliminary analysis showed that not transforming the data provided the best approximation of normally distributed residuals. Thus, growth rate was modeled with a normal distribution and identity link, using the package *lme4* (Bates et al. 2015). The development rate was evaluated by comparing the presence of the different life stages (i.e., larva, pupa, or adult) weighted by the number in each patch, using a cumulative linkage model for ordinal data to consider the order of appearance of each stage (larva, then pupa, then adult), using the package *ordinal* (Christensen, 2015). For the growth rate model and for the development rate model, explanatory variables in the model were the categorical demographic history and temporal blocks as fixed effects, with population identity included as a random effect to account for replication within populations (Table 2). In addition, the observation time (4 weeks or 5 weeks) was also included as a fixed effect in the developmental rate model, as well as the replicate identity as a random effect (Table 2). To consider the variance in population size within each bottleneck category, and thus be able to consider the entire demographic history, we performed these analyses using the expected heterozygosity at the end of the experiment, Het, as a continuous measure of demographic history. We present only the results using the model with the categorical variable for the development stage analysis due to a lack of differences and qualitatively similar findings for both models (see results).

**Table 2.**
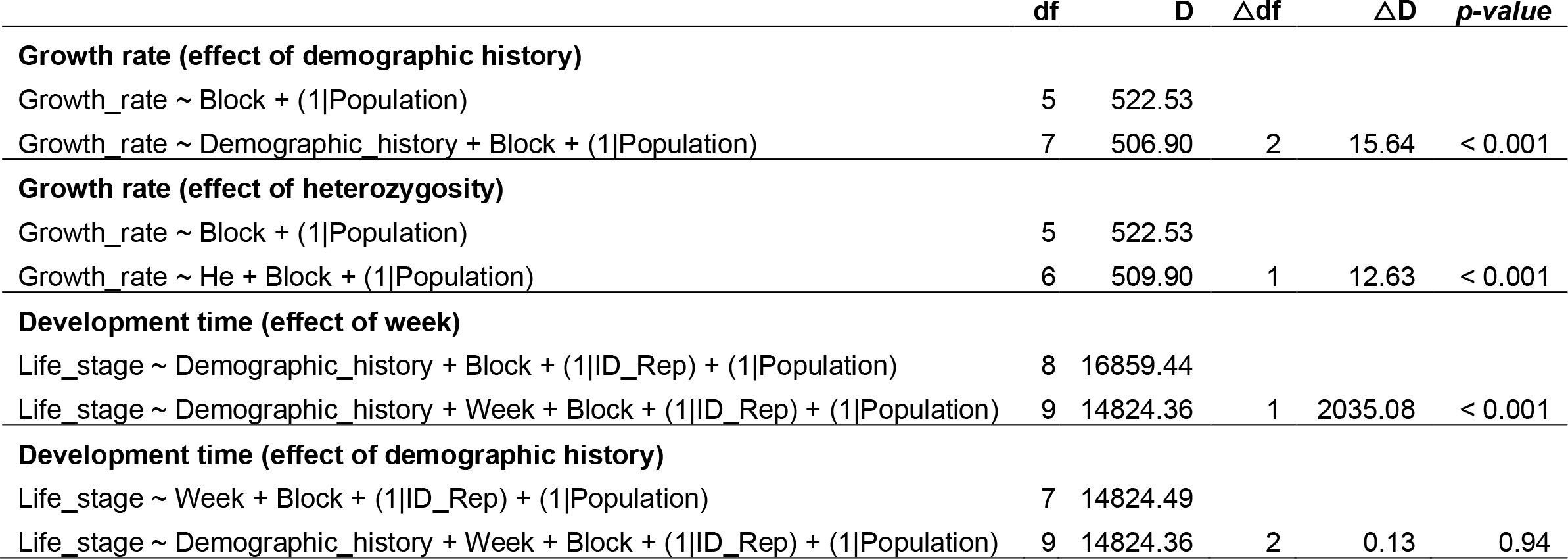
Likelihood ratio test table for each model comparison analyzing the data from assays conducted after the experiment. df: degrees of freedom. D: deviance computed from the likelihood. △df: change in degrees of freedom between the two models. △D: change in deviance between the two models (i.e., statistic of the LRT following a X2 distribution). p-value: p-value of the LRT.

As a complementary step, we estimated the uncertainty of the difference in these traits between strong and intermediate bottlenecks, when the difference was estimated to be small in the analyses described above. To do this, we estimated the 95% confidence interval of the difference on the response scale using a parametric bootstrap assuming the fitted model was the data generating process (5,000 samples, percentile method; Davison and Hinkley 1997). Knowing the confidence interval of the difference allows us to differentiate between the case where the populations are biologically similar (small mean difference with small confidence interval reflecting a narrow range of differences compatible with the data) or if we lack evidence to precisely determine the difference between these two types of bottlenecked populations (small mean difference but large confidence interval reflecting a wide range of differences compatible with the data).

## Results

### Extinction

There was a significant effect of demographic history on the probability of extinction (*P* = 0.009). Populations that had a history of a bottleneck experienced more extinctions in the challenging environment than diverse populations (21.7% and 23.5% of intermediate bottleneck and strong bottleneck populations went extinct, Fig. 2). The absence of bottleneck events during the previous demographic history prevented extinction entirely (Fig. 2, left panel; 0% extinction among diverse populations; estimate of the probability of extinction = 0.004, 95% CI [0.001, 0.05], Table S1). Considering only the demographic histories that experienced extinctions, the probability of extinction was not significantly impacted by bottleneck intensity during demography history (estimate of the probability of extinction for intermediate bottlenecked populations: 0.07, [0.01; 0.27] and for strong bottleneck populations: 0.08, [0.01; 0.28], Table S1, Fig. S1). The difference in probability of extinction between strong and intermediate bottlenecked populations is most likely small, 0.01, but a difference as large as 0.14 is compatible with the data and model assumptions (95% CI of the difference: [-0.12; 0.14]).

**Figure 2.**
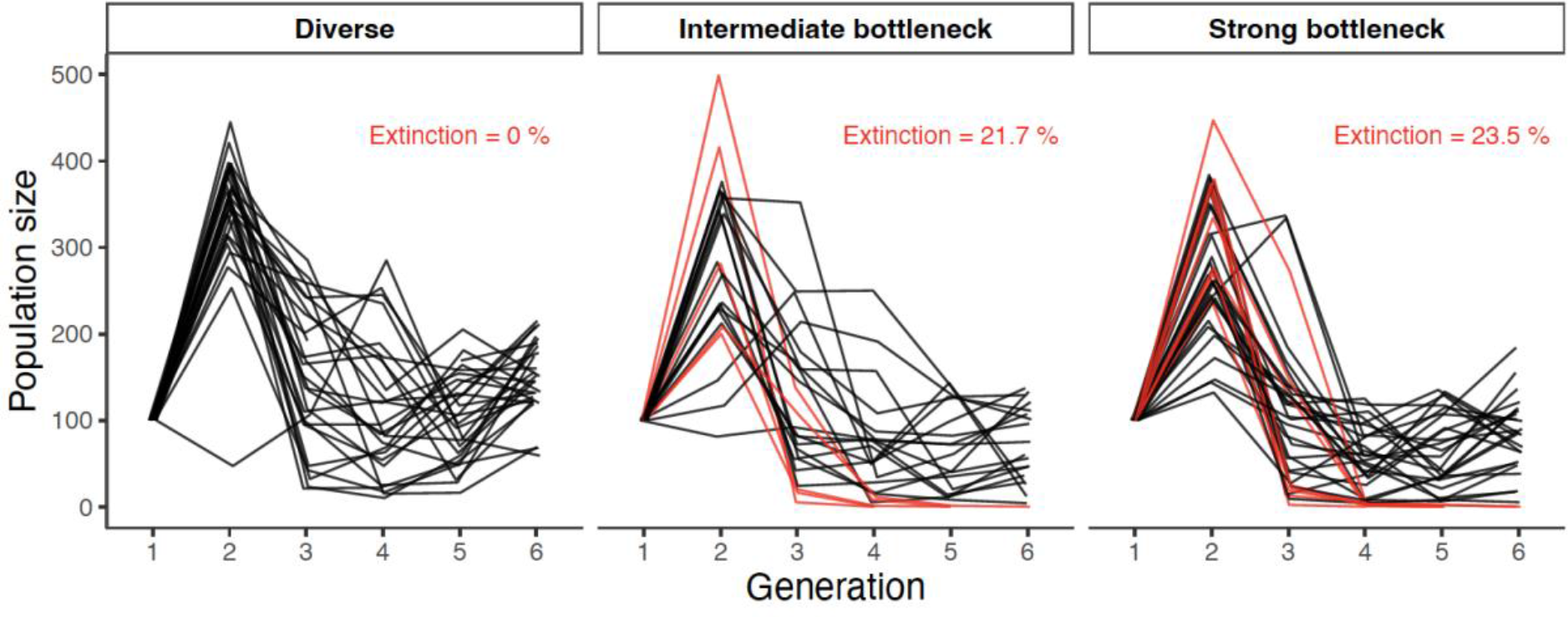
Population size of diverse, intermediate bottleneck, and strong bottleneck populations through time. Red lines indicate populations that went extinct.

Among the populations that went extinct, there was a lack of evidence that the number of generations to extinction differ by demographic history, with largely overlapping confidence intervals (estimates of the time of extinction for intermediate bottlenecked populations: 4.8 generations, [3.1; 7.0] and for strong bottleneck populations: 5.0 generations, [3.6; 6.7] using the model including the demography history effect, Table 1, Table S1). The difference in number of generations to extinction between strong and intermediate bottlenecked populations is most likely small, 0.02, but a difference as large as 2.68 is compatible with the data and model assumptions (95% CI of the difference: [-2.23; 2.68]).

For populations that went extinct, there was not necessarily a gradual decrease in population size that led up to extinction (Fig. S2). Instead, population size mostly declined markedly one generation before extinction (Fig. 2, red-barred circles in Fig. S2), reflecting the sharp population declines.

### Population size dynamics

Following introduction into the challenging novel environment, population sizes initially increased, due to the maternal effects from the high-quality environment the populations were in just prior to the experiment being initiated (as shown by Van Allen and Rudolf 2013), and then dropped dramatically (Fig. 2).

In extant populations, analysis of population size over time reveals U-shaped demographic curves indicative of evolutionary rescue (Fig. 3). The estimated curves from the best-fit GAMM model, where the shapes are the same regardless of the demographic history (the most parsimonious model), are shown in Fig. 3A. The estimated curves from the full GAMM model, allowing different shapes depending on the demographic history (best representing the experimental design), are shown in Fig. 3B (Table S2). There are not statistically clear differences in the curves by demographic history (interaction between demographic history and generation allowing different curves: *P* = 0.42, Table 1). Although the overall U-shape was apparent for all treatments, mean population size was larger for the diverse- population treatment than for the intermediate-bottleneck and strong-bottleneck treatments (demographic history effect: *P* < 0.001, Table 1; confidence interval for the diverse populations does not overlap in Fig. 3A or diverges in Fig. 3B from intermediate and strong bottleneck populations). The predicted mean population size of the intermediate-bottleneck populations was slightly higher than for strong- bottleneck populations, but confidence intervals overlapped for all generations (Fig. 3, Table S2). By the last generation, the mean population size of the extant populations was greater than the initial population size only for the diverse populations (confidence interval not including 100 for the diverse demography history; Figs. 2, 3). Indeed, by the last generation, the diverse populations had reached a mean population size of 127.9 (95% CI [107.3, 152.4]) individuals, while the intermediate bottleneck and strong bottleneck populations had reached means of 86.8 [71.3, 105.7] individuals and 80.4 [67.3, 96.0] individuals respectively, according to the estimates from the best-fit GAMM model (lines in Fig. 3A). These estimates remain similar when using the full model that best represents the experimental design (population size at the last generation for diverse populations: 143.5 [116.3, 177.2], for intermediate bottleneck: 74.6 [58.0, 96.0] and for strong bottleneck: 76.8 [61.6, 95.6]; Fig 3B). The estimated mean of the extant diverse populations from the best-fit GAMM model was always either above or overlapping the initial size (minimum of 96.0 individuals at 4.7 generations but with a confidence interval including 100 individuals). This was not the case for the bottlenecked populations where the population size fell to a mean of 65.2 individuals and 60.3 individuals for the intermediate bottleneck and strong bottleneck respectively (confidence interval not including 100 for generations 4 and 5 in Fig. 3A). These predictions remain similar when using the full model that best represents the experimental design (minimum of the population size for diverse populations: 99.8 [84.7, 117.6], for intermediate bottleneck: 62.0 [50.7, 75.8] and for strong bottleneck: 58.4 [49.0, 69.5]; Fig 3B).

**Figure 3.**
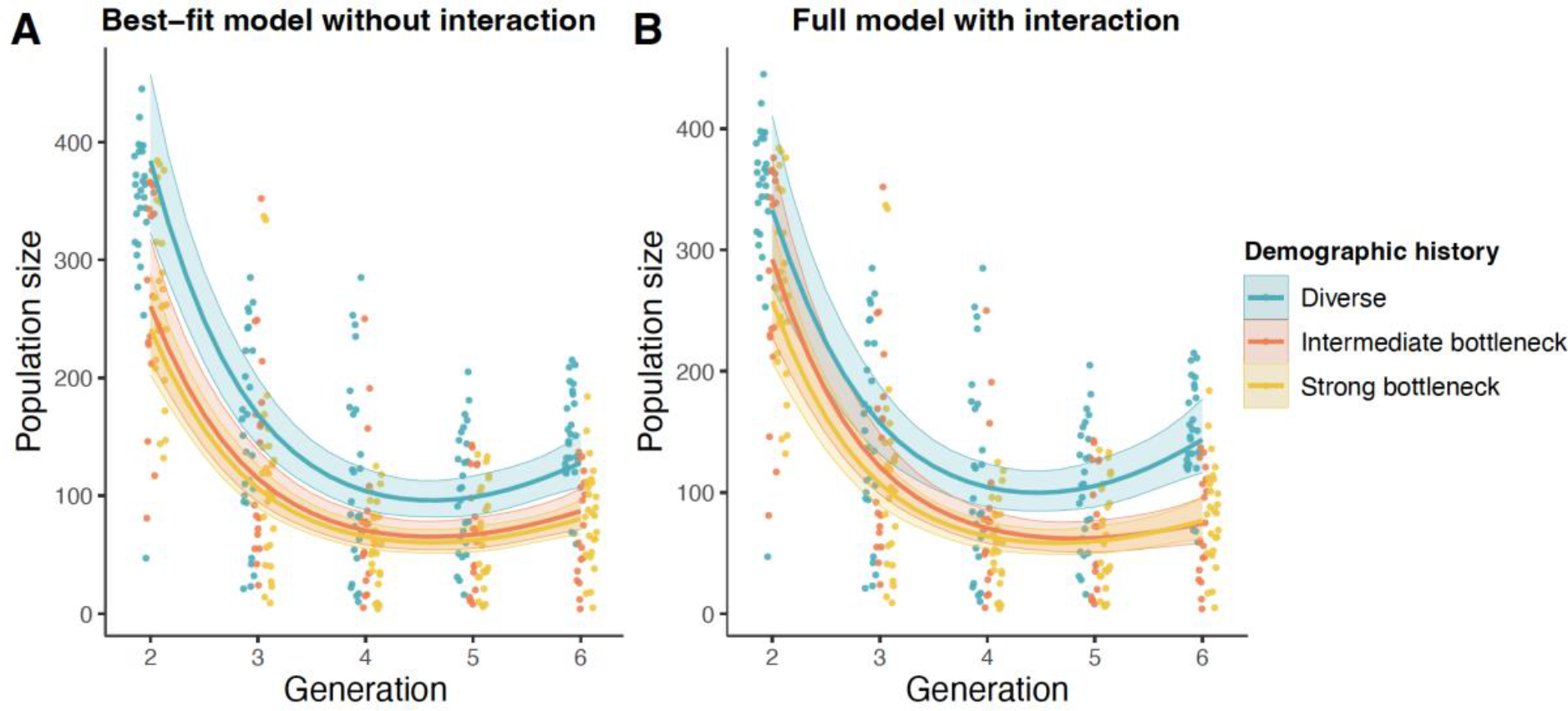
Population size of extant diverse populations (blue), intermediate bottleneck populations (red) and strong bottleneck populations (yellow) through time estimated by (A) the best-fit GAMM model or (B) the full GAMM model. Each population is represented by a point. The lines are estimates of mean population size for the three demographic history levels from the GAMM model, averaged over the three temporal blocks. Data include extant populations beginning at generation 2. The best-fit GAMM model does not include an interaction between the demographic history and the time, which means that the shape of the predictions cannot differ by demographic history. The full GAMM model includes an interaction between demographic history and time, which means that the shape of the curve can differ by demographic history. The shaded and colored area around each line is the 95% confidence interval. Points in different treatments are staggered slightly and jittered on the generation axis to reduce overlap (generation is discrete).

Coefficients of variation for population size across the three demographic histories for almost all generations had overlapping confidence intervals (Fig 4). For most generations, diverse populations had the lowest coefficient of variation, and at generation 6, population sizes of the extant diverse populations were less variable than the extant intermediate bottleneck and strong bottleneck populations (CVdiverse = 0.29 [0.22, 0.41]; CVintermediate = 0.60 [0.42, 1.07], CVstrong = 0.51 [0.38, 0.77]; Fig. 4).

**Figure 4.**
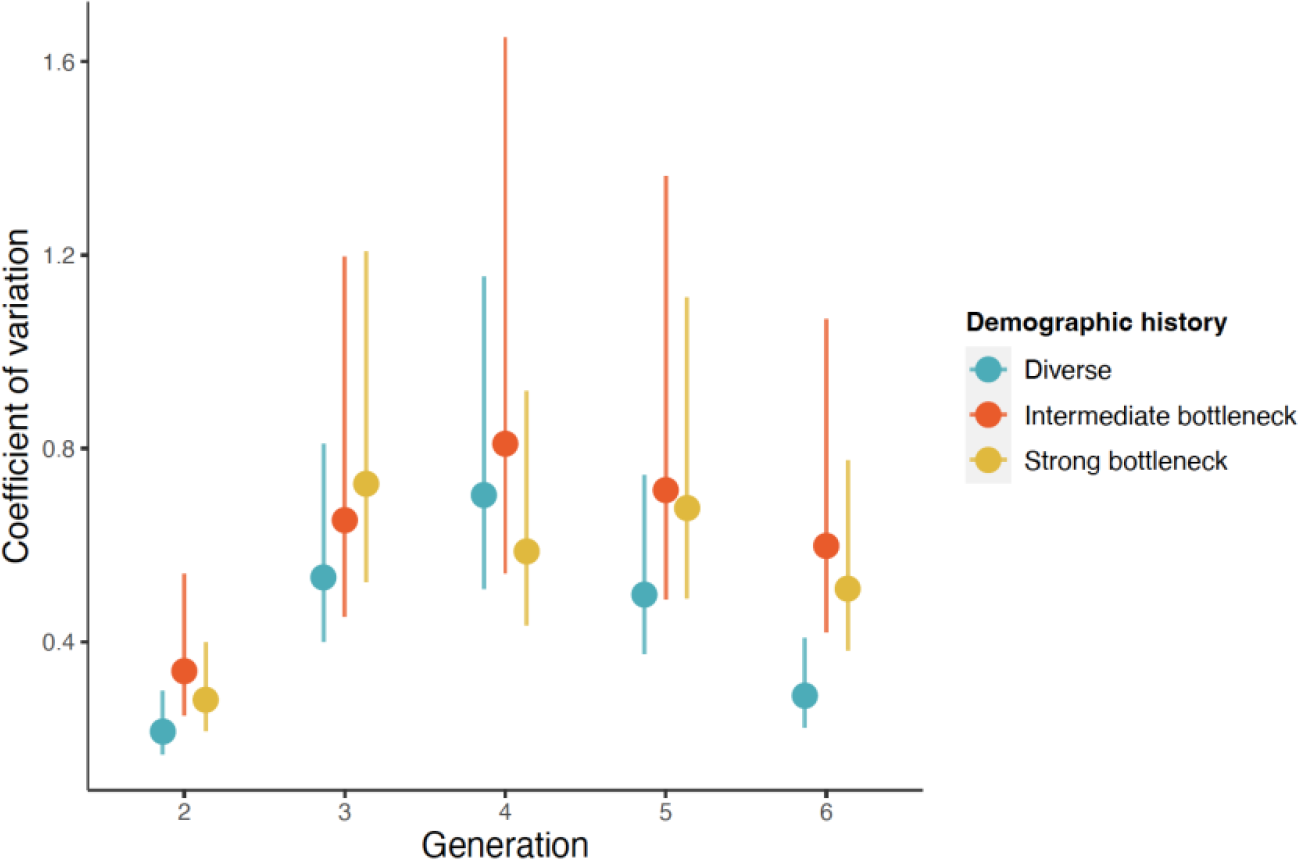
Coefficient of variation of population size through time. The mean coefficient of variation for each demographic history is represented by a point, with the 95% confidence interval represented by an error bar. Extinct populations were excluded but results were similar when they are considered. Points in different treatments are staggered slightly on the generation axis to reduce overlap (generation is discrete).

These results remained unchanged even when considering all populations, including the populations that went extinct as having a population size of zero (results not shown).

### Growth rate and development time

Most of the extant populations for the three demographic histories had on average at the end of the experiment, a growth rate higher than 1, which means that more descendants were produced than there were parents (Fig. 5). At the end of the experiment, the growth rate was significantly different depending on the demographic history (demography history effect: *P* < 0.001, Table 2). The growth rate was higher for the diverse populations (estimate: 2.57, 95% CI [2.20, 2.94], Table S3), than for bottlenecked populations (estimate: 1.70 [1.26, 2.13] for intermediate and 1.87 [1.50, 2.23] for strong bottleneck populations, Table S3). The difference in growth rate between strong and intermediate bottlenecked populations is most likely small, -0.17, but a difference of up to -0.69 is also consistent with the data and model assumptions (95% CI of the difference: [-0.67; 0.35]). Overall, the growth rate increased significantly with the heterozygosity of the populations (slope estimate: 1.77 [0.85, 2.68]; *P* < 0.001, Table 2).

**Figure 5.**
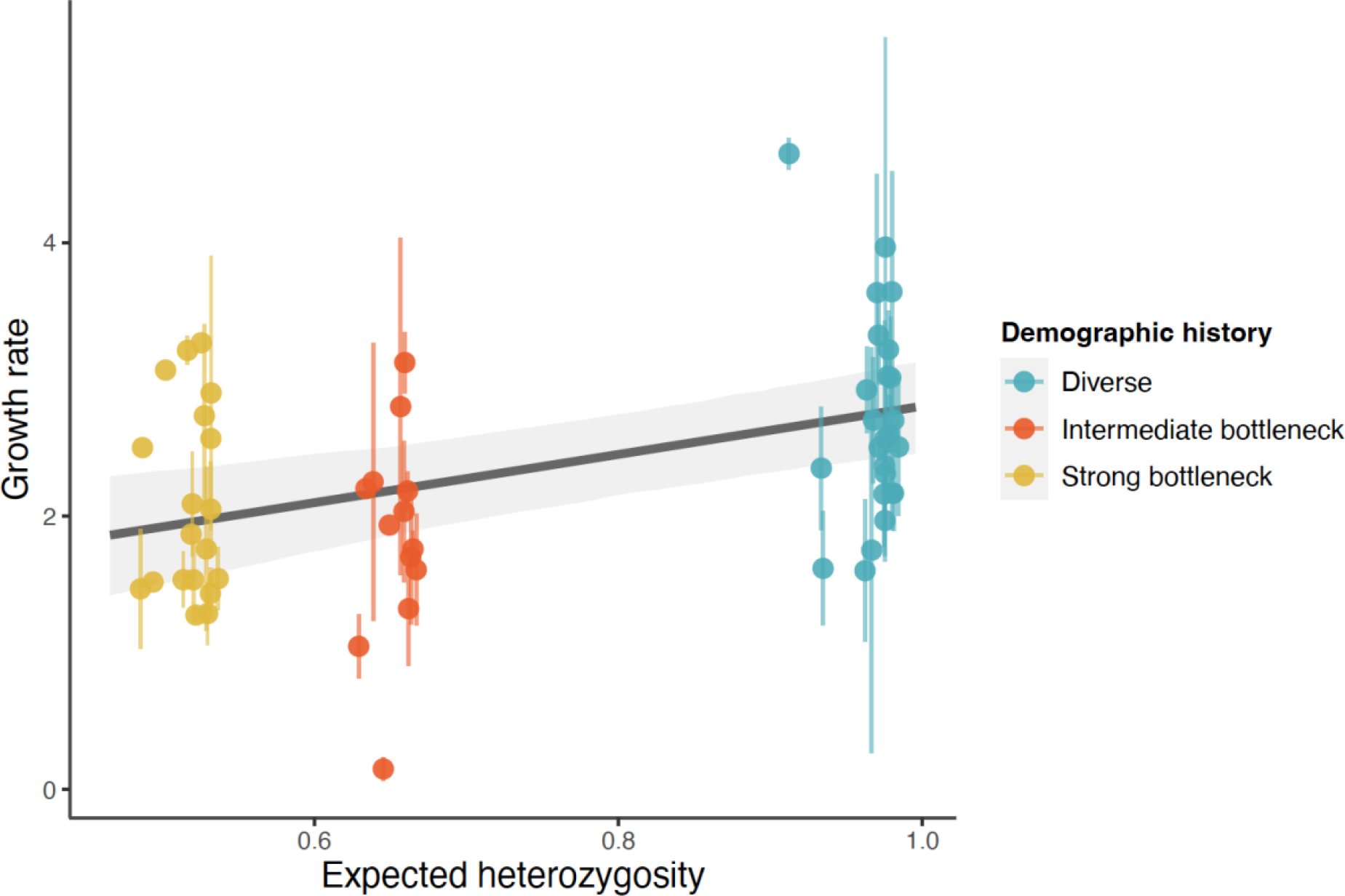
Growth rate assays for each extant population after the rescue experiment depending on its final heterozygosity. The mean growth rate across assays of each population is represented by a point, with the standard error represented by an error bar. The points without error bars are from populations where there was only one replicate assay to estimate the mean. The solid gray line is the estimated mean relationship from the linear model. The shaded area is the 95 percent confidence interval.

The demographic history of rescued populations had little to no influence on developmental timing. The proportions of larvae, pupae, and adults were similar among the three demographic histories (Fig. S3) and there was insufficient evidence for a difference (*P* = 0.94, Table 2). These proportions varied significantly only with respect to timestep, i.e., between 4 weeks and 5 weeks (*P* < 0.001, Table 2).

## Discussion

In this study, we evaluated how the demographic history of populations influences the probability of extinction and adaptation to a challenging environment. We found that populations that experienced a bottleneck event in their demographic history were more likely to go extinct, and, if they survived, had smaller population sizes, and were less well adapted to the challenging environment than populations that had not experienced bottleneck events.

### Extinction associated with large initial drop in population size for bottlenecked **populations**

While all diverse populations persisted throughout the six generations of the experiment, multiple bottlenecked populations went extinct. These extinctions were probably due to the high inbreeding depression and loss of genetic diversity present in bottlenecked populations (Frankham et al. 2019). Notably, except for one population, none of the populations that experienced an initial drastic drop in population size associated with a 55% reduction heterozygosity, were able to avoid extinction. This may be due to an extinction vortex, a process in which small populations remain small due to a reinforcing negative feedback loop that makes them vulnerable to extinction due to demographic stochasticity (Gilpin & Soulé, 1986). The diverse populations did not decline to as small a size as the bottlenecked populations, and perhaps this enabled them to avoid falling into an extinction vortex. In the future, it would be interesting to know if diverse populations that decline more sharply, due to stochasticity or experimental manipulation, can escape more easily via rapid adaptation than populations that have experienced bottleneck events.

### Dynamics of rescue for extant populations

Extant populations from all treatments exhibited the U-shaped population size trajectory characteristic of evolutionary rescue. Diverse populations showed a higher level of adaptation, quantified by population growth rate, than the bottlenecked populations. These results confirm that rapid adaptation is compromised following a bottleneck event (Bouzat, 2010). Our findings are consistent with those of Klerks *et al*., (2019) who showed that bottlenecked populations had a 50% slower response to selection for an increased heat tolerance than normal populations. Further investigation should consider whether this difference is due to a reduction of genetic variation in bottlenecked populations or an increase of inbreeding depression, or both. This would require a combination of phenotypic and genetic analysis, and could help would facilitate efficient management of populations where genetically variable and unrelated individuals are often not available. For instance, it would increase the probability of success of a translocation effort by targeting the individuals to be introduced (Ralls *et al*., 2020). If the effects of bottlenecks are primarily driven by a lack of genetic variation, individuals evolving in different environments, even if they have a recent common ancestor, should be considered for introductions to natural populations, to maximize the probability of having a large pool of different alleles. However, if the effects of bottlenecks are primarily driven by inbreeding depression, then the introduction of unrelated individuals, even if they are all adapted to the same environment with all the adaptive alleles to their environment potentially fixed and thus lacking genetic variation in those traits, could be more beneficial.

### Similar patterns of evolutionary rescue for intermediate and strong bottleneck **populations.**

The evolutionary responses in terms of extinction, rescue, adaptive dynamics, and traits involved were similar for populations that experienced intermediate bottlenecks and populations that experienced strong bottlenecks. It is important to note that a small difference may exist, but our experimental design does not have the power to detect it. Indeed, while the difference was always small, whatever the phenotype, the upper limit of the confidence interval of this difference was large for most phenotypes. However, if a biological difference between intermediate and strong bottlenecks exists, it must be smaller than that observed between bottleneck and diverse populations. Thus, here we discuss three alternative hypotheses to explain the general congruence between the two bottleneck treatments. First, the difference in expected heterozygosity between intermediate and strong bottleneck events at the start of the experiment was smaller than that between bottlenecked and diverse populations (see Fig. S1; averages at the start of the experiment: Hediverse = 0.99, Heintermediate = 0.68, Hestrong = 0.54). Both types of bottleneck reduced heterozygosity substantially, but still some heterozygosity remained. It may be that in this range of expected remaining heterozygosity, evolutionary responses to the environment are similar. Second, populations that experienced a strong bottleneck event may have purged deleterious alleles (Kirkpatrick & Jarne, 2000), reducing the negative fitness effects of the bottleneck and making phenotypic outcomes more similar to the intermediate bottleneck. Although many studies have failed to find clear evidence, the process of purging has been demonstrated in some natural populations (Bouzat et al. 2010, Bertorelle et al. 2022). For instance, a recent analysis of the genomes of wild populations of Alpine ibex that had undergone severe bottlenecks revealed a purge of highly deleterious alleles, demonstrating that in certain circumstances purging can be partially effective (Grossen *et al*., 2020). In our study, the level of adaptation to the challenging environment may be comparable in intermediate and strong bottlenecks, despite differences in heterozygosity, due to potential purging of deleterious alleles in strong bottlenecks. This potential purge could be tested by comparing the fitness of offspring from inbred or outbred crossing (e.g., Facon et al. 2011) in each of these populations. Third, even if the phenotypic patterns seem similar, the adaptive pathways used by these two types of populations may be different. For instance, Vogwill *et al*., (2016) found that antibiotic resistance evolved through highly beneficial mutations for bacterial populations that experienced a strong bottleneck event, whereas it evolved through a greater diversity of genetic mechanisms in populations that experienced an intermediate bottleneck event.

Differences in the adaptive pathways may have consequences on the future evolutionary potential of these populations.

### Variance of evolutionary trajectories may depend on the demographic history **of populations.**

Evolutionary responses were more variable in strong and intermediate bottlenecked populations than in diverse populations, but the confidence intervals overlapped throughout the experiment, except for the last generation. Theoretical expectations predict higher variance for bottlenecked populations due to sampling of the genetic composition of each replicate population during a bottleneck event (Bouzat, 2010). The lack of support for differing coefficients of variation at the beginning of the experiments might be because all populations were introduced into a new environment and experience it in a similar way. In addition, contrary to theoretical predictions and experimental results (Vogwill *et al*., 2016; but see Windels *et al*., 2021), we did not observe significantly higher coefficients of variation for strong bottlenecks than for intermediate bottlenecks. As described above, this lack of evidence for a difference could be due to a lack of statistical power or a lack of difference due to purging or a lack of phenotypic difference (see the section on similar responses between bottlenecked populations for more details).

### Life history traits involved in the adaptation process

To better understand the different levels of adaptation between treatments, we examined the proportions of individuals in different life stages at two time points at the end of the experiment, to search for potential differences in development time. We found that these proportions were on average very similar among demographic histories. Life history traits other than development time may thus be responsible for adaptation to the challenging environment. For instance, Agashe *et al*., (2011) demonstrated an increase of fecundity and egg survival, as well as a higher rate of egg cannibalism, associated with adaptation to a new corn environment for populations of *T. castaneum*.

### Contributions and ideas for future genomic research projects

While genomic analyses were not within the scope of this study, this project opens up many fruitful avenues of research, as mentioned above, that could be addressed with genomics. We propose three concrete ways in which sequencing these types of experimental or natural populations would address the different perspectives mentioned above. First, sequencing these types of populations would allow us to measure the components of genetic load (i.e., “the proportion by which the population fitness is decreased in comparison with an optimum genotype”) as described in Bertorelle et al. (2022) to determine whether genetic load can be used as good indicator for predicting extinction probabilities. Secondly, sequencing populations whose population dynamics are known would allow us to directly measure and calculate heterozygosity to know if this measure allows us to correctly describe the demographic history of populations (Frankham 2002, Frankham 2010). Third, by looking at populations adapted to different stressors, it would be interesting to know if the pathways used are more diverse involving a large range of polygenic adaptation, when the populations have or have not undergone complex demographic histories (Gamblin et al. 2023, Garoff et al. 2020).

### Conclusion and perspectives

Our findings have implications for conservation in today’s changing world. Managers often have been concerned with maintaining adaptation to local environments for populations of conservation concern (Dalrymple et al., 2021). In the Anthropocene, where environments are changing extremely rapidly, a focus on supporting sufficient genetic variation for ongoing and future adaptation to shifting conditions is important. The reduced ability of bottlenecked populations in this study to adapt to altered conditions suggests that more active management strategies may be needed for populations that have experienced bottlenecks. Therefore, our work supports the substantial literature on genetic rescue, the process of alleviating inbreeding depression and restoring genetic diversity of inbred populations through gene flow (Whiteley et al. 2015, Ralls *et al*., 2020).

### Data Archiving Statement

Data for this study are available at: to be completed after manuscript is accepted for publication. The data and R scripts for our analyses are available at: https://anonymous.4open.science/r/AnalysisEvolRescueExp-4051/

## Acknowledgments

L.O., B.L. and R.A.H. acknowledges support from the department Agricultural Biology at Colorado State University for the Ag Biology Undergraduate Research Fellowship. L.O., R.A.H. and B.A.M. acknowledge support from the US National Science Foundation (DEB-1930650 to R.A.H.; DEB-1930222 to B.A.M.).

