## Supplementary materials for "Population demographic history and evolutionary rescue: influence of a bottleneck event"

**Running Title (max 40 characters):**
Demographic history and evolutionary rescue

### Supplementary tables

***Table S1. Estimates for the probability of extinction model and the time to extinction model.*** *The estimates for the probability of extinction model represent the best model (LRT model selection). For the time to extinction, this model is not the best model since the demographic history effect was not significant. Estimates were back-transformed to facilitate biological interpretation. The 95% confidence interval of each parameter is reported with the lower and upper confidence limits (CI_2.5% and CI_97.5% respectively).*

| **Extinction_probability ~ Demographic_history + Block** | **Estimates** | **CI_2.5%** | **CI_97.5%** |
|---|---|---|---|
| Demographic_history:Diverse | 0.004 | 0.00002 | 0.05 |
| Demographic_history:Intermediate | 0.07 | 0.01 | 0.27 |
| Demographic_history:Strong | 0.08 | 0.01 | 0.28 |
| Block:Block4 | 0.63 | 0.17 | 0.95 |
| Block:Block5 | 0.93 | 0.71 | 0.99 |
| **Time_extinction ~ Demographic_history** | **Estimates** | **CI_2.5%** | **CI_97.5%** |
| Demographic_history:Intermediate | 4.80 | 3.13 | 6.98 |
| Demographic_history:Strong | 5.00 | 3.61 | 6.71 |

***Table S2. Estimates for the population size model.*** *This GAMM model is the full model but is not the best model since the interaction between demographic history and the generation effect (represented in the model with the s(Generation, by = Demographic_history) term) was not significant. The population size expected with this model and these estimates are shown in Figure 3B.*

| **Population_size ~ Demographic_history + Block + s(Generation) + s(Population, bs="re") + s(Generation, by = Demographic_history)** | | |
|---|---|---|
| *Parametric effects:* | **Estimates** | **Std.error** |
| Demographic_history:Diverse | 5.17 | 0.08458 |
| Demographic_history:Intermediate | 4.76 | 0.08912 |
| Demographic_history:Strong | 4.65 | 0.07734 |
| Block:Block4 | -0.09 | 0.08693 |
| Block:Block5 | -0.34 | 0.10114 |
| *Smooths effects:* | **edf** | **ref.df** |
| s(Generation_Eggs) | 3.37 | 3.78 |
| s(Generation_Eggs):Diverse | 1.00 | 1.00 |
| s(Generation_Eggs):Intermediate | 0.01 | 0.01 |
| s(Generation_Eggs):Strong | 1.00 | 1.00 |
| s(Population) | 19.88 | 66.00 |

***Table S3. Estimates for the growth rate models and the development time model.*** *For each model, the model shown corresponds to the best model (LRT model selection). For the fixed effects, the 95% confidence interval of each parameter is reported with the lower and upper confidence limits (CI_2.5% and CI_97.5% respectively). For the random effects, the variance is reported with its standard deviation.*

| | | | |
|---|---|---|---|
| **Growth_rate ~ Demographic_history + Block + (1\|Population)** | | | |
| ***Fixed effects:*** | **Estimates** | **CI_2.5%** | **CI_97.5%** |
| Demographic_history:Diverse | 2.57 | 2.20 | 2.94 |
| Demographic_history:Intermediate | 1.70 | 1.26 | 2.13 |
| Demographic_history:Strong | 1.87 | 1.50 | 2.23 |
| Block:Block4 | 0.22 | -0.19 | 0.63 |
| Block:Block5 | 0.01 | -0.48 | 0.47 |
| ***Random effects:*** | **Variance** | **Std.dev** | |
| Population | 0.23 | 0.48 | |
| Residuals | 0.75 | 0.86 | |
| **Growth_rate ~ He + Block + (1\|Population)** | | | |
| ***Fixed effects:*** | **Estimates** | **CI_2.5%** | **CI_97.5%** |
| Intercept | 0.82 | 0.10 | 1.53 |
| He | 1.77 | 0.85 | 2.68 |
| Block:Block4 | 0.22 | -0.20 | 0.64 |
| Block:Block5 | -0.01 | -0.49 | 0.48 |
| ***Random effects:*** | **Variance** | **Std.dev** | |
| Population | 0.25 | 0.50 | |
| Residuals | 0.75 | 0.87 | |
| **Life_stage ~ Week + Block + (1\|ID_Rep) + (1\|Population)** | | | |
| ***Fixed effects:*** | **Estimates** | **CI_2.5%** | **CI_97.5%** |
| Larvae\|Pupae | -5.31 | -5.66 | -4.97 |
| Pupae\|Adults | -3.42 | -3.75 | -3.09 |
| Week:Week4 | -2.62 | -2.75 | -2.48 |
| Block:Block4 | 0.10 | -0.34 | 0.55 |
| Block:Block5 | 0.17 | -0.38 | 0.72 |
| ***Random effects:*** | **Variance** | **Std.dev** | |
| ID_Rep | 0.52 | 0.72 | |
| Population | 0.34 | 0.58 | |

### Supplementary figures

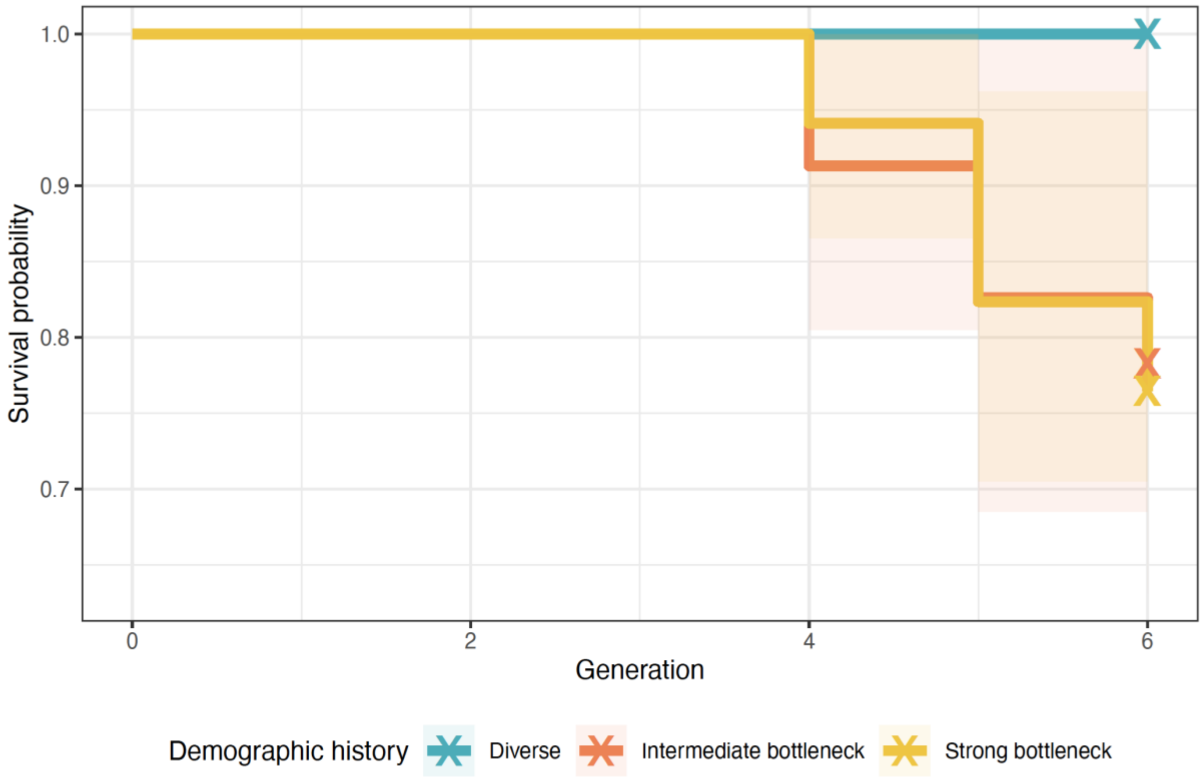


***Figure S1. Survival probability through time for each demographic history.*** *The lines represent the model estimate of overall survival for the three demographic history levels predicted by a survival model: Surv(time_to_extinction, status_extinction) ~ Demographic_history, using the function surv() from the package survival(). The shaded and colored area around each line represents the 95% confidence interval. The crosses represent the last censored observations (i.e., the experiment). Survival curves differed across demographic histories ($\chi^2_2$ = 7, P = 0.03) using a log-rank test with the survdiff() function.*

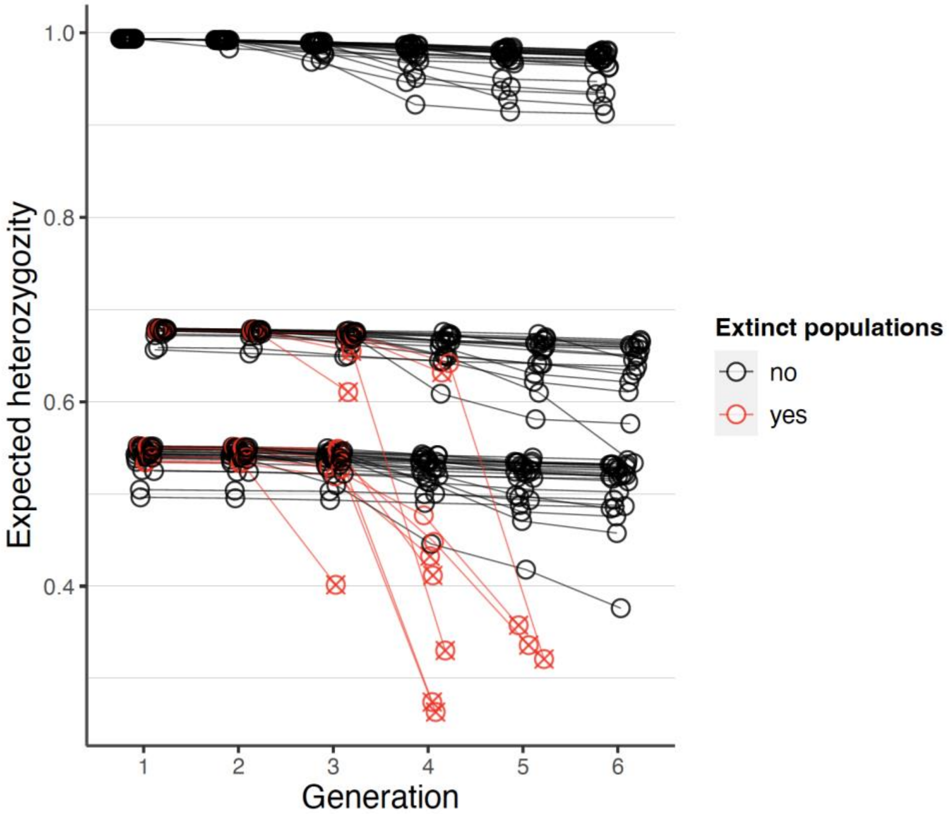


***Figure S2. Expected heterozygosity through time.*** *Red lines indicate populations that went extinct. For extinct populations, the barred circle represents the expected heterozygosity at the last generation before extinction. Points are jittered on the generation axis to reduce overlap.*

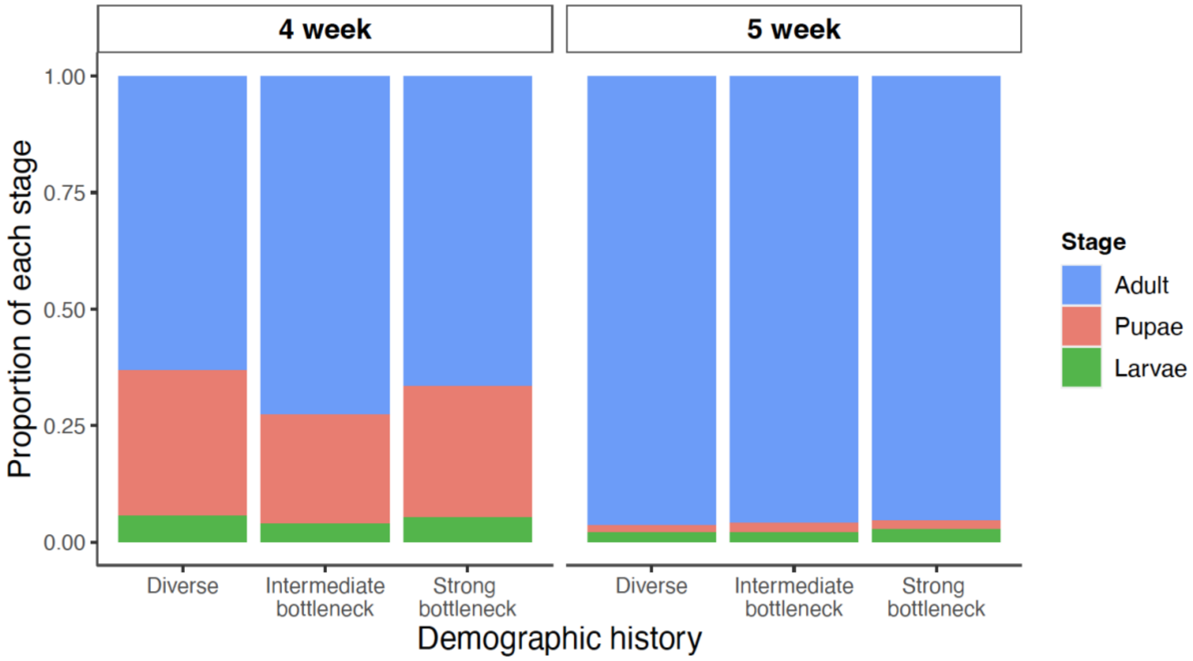


***Figure S3. Proportion of each life-stage after 4 weeks (left panel) and 5 weeks (right panel) of development time, for the three demographic history treatments.***
